## Supplementary material for "Nanopore-Based Enrichment of Antimicrobial Resistance Genes – A Case-Based Study": SFig. 1-3

547 # Supplement

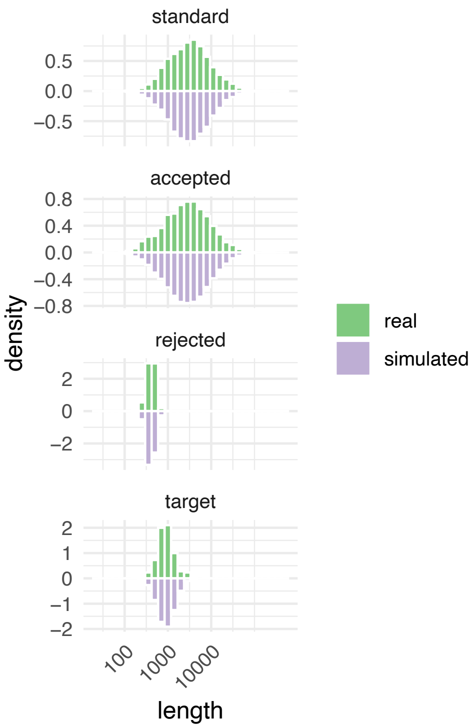


Figure S1: Exemplary length distribution of various sequence sets of interest discussed in the study (isolate A2). Generated reads and target genes in green, simulated sequences in violet. Note that the x-axis is on the log scale. As expected, reads generated using standard sequencing do not differ in their length distribution from unrejected reads from adaptive sampling. Only the sequences of rejected reads are truncated, usually after on average 415 bases (median). The antimicrobial resistance genes in our target database have a mean length of 1010 bases. All displayed distributions can be modeled using a log-normal distribution. By varying model parameters, we can simulate unobserved, counterfactual read-target combinations. We then use these pairs to assess the false-negative rate of target detection (see results). Actual and simulated sequence length distributions are near-identical, suggesting that the results derived from the simulation are realistic.

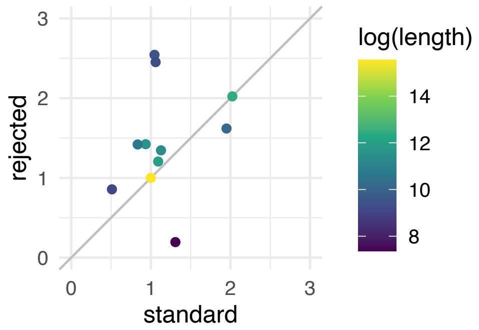


Figure S2: Relative coverage of reads generated using standard sequencing and the rejected read fraction from adaptive sequencing (isolate A2). Adaptive sequencing has been shown not to change the relative abundance of reads *outside of target regions*.[12] Correspondingly, we expect the same coverage from both groups, which we find, validating the correctness of the adaptive sequencing run. Note that we normalized the coverage of all contigs within each condition to the coverage of the chromosome because rejected reads are more numerous and shorter. Comparing unadjusted coverages would artificially inflate coverages from the rejected reads. The normalized coverage is similar between standard and rejected reads, except for the smallest contigs. Here we see a slight deviation, which due to the small contig size does not affect the conclusions in this study.

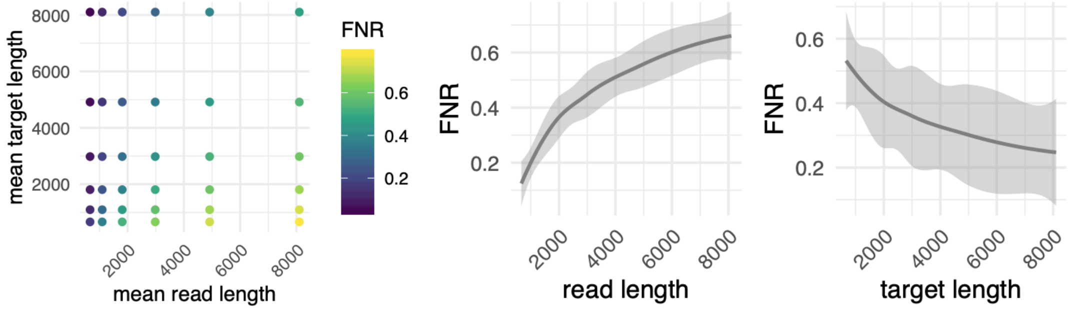


Figure S3: Simulation of counterfactual read-target pairs for various length distributions shows a large effect on the number of false-negative read rejections. The shorter a target relative to the median read length of the sequencing run, the larger the false-negative rate (FNR, left panel, see results). For the use case discussed in this manuscript, namely the highly multiplexed detection of antimicrobial resistance genes, it is beneficial then the median read length matches the target size. A multivariate regression on the simulated pairs estimates this effect in further detail (middle and right panel).

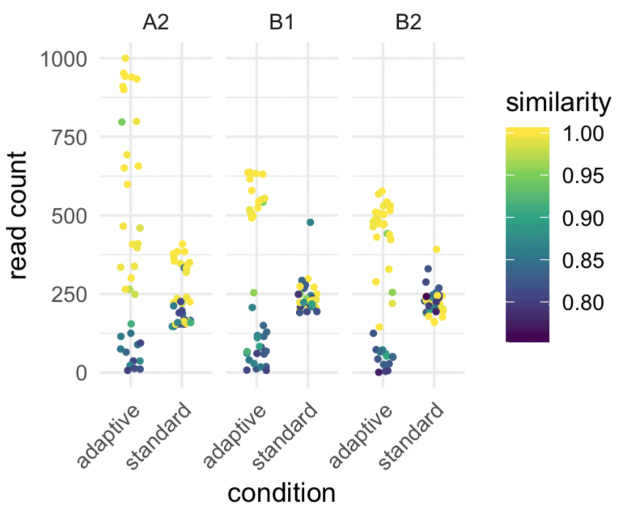


Figure S4: While the main manuscript analyses the pooled reads from three isolates, we here demonstrate that the effects observed in aggregate reproduce at the level of the individual isolates (compare Figure 2B).
